# Learning with phenotypic similarity improves the prediction of functional effects of missense variants in voltage-gated sodium channels

**DOI:** 10.1101/2022.09.29.510111

**Authors:** Christian Malte Boßelmann, Ulrike B.S. Hedrich, Holger Lerche, Nico Pfeifer

## Abstract

**Background:** Missense variants in genes encoding voltage-gated sodium channels are associated with a spectrum of severe diseases affecting neuronal and muscle cells, the so-called sodium channelopathies. Variant effects on the biophysical function of the channel correlate with clinical features and can in most cases be categorized as an overall gain- or loss-of-function. This information enables a timely diagnosis, facilitates precision therapy, and guides prognosis. Machine learning models may be able to rapidly generate supporting evidence by predicting variant functional effects.

**Methods:** Here, we describe a novel multi-task multi-kernel learning framework capable of harmonizing functional results and structural information with clinical phenotypes. We included 62 sequence- and structure-based features such as amino acid physiochemical properties, substitution radicality, conservation, protein-protein interaction sites, expert annotation, and others. We harmonized phenotypes as human phenotype ontology (HPO) terms, and compared different measures of phenotypic similarity under simulated sparsity or noise. The final model was trained on whole-cell patch-clamp recordings of 375 unique non-synonymous missense variants each expressed in mammalian cells.

**Results:** Our gain- or loss-of-function classifier outperformed both conventional baseline and state-of-the-art methods on internal validation (mean accuracy 0.837 ± 0.035, mean AU-ROC 0.890 ± 0.023) and on an independent set of recently described variants (n = 30, accuracy 0.967, AU-ROC 1.000). Model performance was robust across different phenotypic similarity measures and largely insensitive to phenotypic noise or sparsity. Localized multi-kernel learning offered biological insight and interpretability by highlighting channels with implicit genotype-phenotype correlations or latent task similarity for downstream analysis.

**Conclusions:** Learning with phenotypic similarity makes efficient use of clinical information to enable accurate and robust prediction of variant functional effects. Our framework extends the use of human phenotype ontology terms towards kernel-based methods in machine learning. Training data, pre-trained models, and a web-based graphical user interface for the model are publicly available.

## Background

Variants in genes encoding voltage-gated sodium channels are associated with a broad spectrum of often severe neurodevelopmental, neuromuscular, and cardiac diseases. Complementary genetic and electrophysiological investigations have contributed towards a growing list of these ion channel disorders or ‘channelopathies’ (1). For example, variants in sodium channel protein type 1 subunit alpha (*SCN1A*, Na_V_1.1) are the second-most common finding in cohorts of individuals with monogenic epilepsy (2). The majority of these variants are associated with Dravet syndrome, a severe developmental and epileptic encephalopathy (DEE) and the most common drug-resistant genetic epilepsy (3). Likewise, variants in *SCN5A* are responsible for >75% of cases with genetically confirmed Brugada syndrome (BrS), an abnormality of cardiac conduction with a significant risk of sudden death in young individuals (4). Lastly, variants in *SCN9A* are associated with paroxysmal extreme pain disorder (PEPD), a rare autosomal-dominant disorder of pain processing and autonomic function with neonatal to infantile onset, where individuals experience sudden attacks of excruciating pain with no response to conventional analgesic medication (5). These examples serve to illustrate the high burden of disease and unmet medical need in the affected individuals.

Missense variants can have an impact on the biophysical function of the mutant ion channel. Amino acid substitutions may alter the physiochemical properties including polarity, residue weight and size, disulphide linkage, or hydrophobicity of residues that are located in domains or functional motifs that are critical to channel function (6-8). This may affect every aspect of channel function including voltage sensing, expression level or membrane trafficking, all parameters of channel gating such as the voltage-dependence or time constants of activation and fast or slow inactivation, incomplete channel inactivation leading to persistent sodium current, or loss of ion selectivity (9-12). In the majority of cases, the sum of the functional consequences of a variant can be categorized as an overall net gain-of-function (GOF) or loss-of-function (LOF) (13). Less often, mixed effects on channel function, downstream effects on neuronal firing or network dysfunction, or different functional results depending on the expression system contribute towards a variant for which an overall GOF or LOF may be difficult to determine (14, 15).

These functional effects of missense variants correlate with distinct clinical phenotypes, which have been described for almost all voltage-gated sodium channels. While *SCN1A*-LOF is associated with Dravet syndrome, *SCN1A*-GOF causes familial hemiplegic migraine type 3 (FHM3) or a spectrum of DEE including movement disorder and arthrogryposis (NDEEMA) (16, 17). In *SCN5A*, LOF by altered channel inactivation or non-conducting channels leads to Brugada syndrome type 1, while GOF by increased persistent inward sodium current and longer action potential duration results in familial long QT syndrome type 3 (LQT3) (18-21). Perhaps most strikingly, *SCN9A*-LOF causes paroxysmal extreme pain disorder, yet *SCN9A*-GOF leads to congenital insensitivity to pain (22). For *SCN2A* and *SCN8A*, GOF is associated with an early-onset epilepsy phenotype including self-limited (familial) infantile epilepsy (SeLIE) or DEE. Individuals with these variants respond well to treatment with sodium channel blockers (SCBs). Conversely, *SCN2A-*LOF or *SCN8A*-LOF leads to a later seizure onset and unfavourable response to SCBs (23, 24). Evidently, individual phenotypic features contain valuable information to guide variant interpretation.

Knowledge of variant functional effects and genotype-phenotype correlations facilitates a timely diagnosis, informs genetic counselling, and offers therapeutic and prognostic guidance to the clinician (25). However, variant interpretation requires causal evidence generated by electrophysiological experiments which are relatively time-consuming and expensive. Recent advances in next-generation sequencing techniques have generated a wealth of genetic variation which threatens to outpace the ability of experimental assays to keep up with the increased demand, resulting in a growing number of individuals affected by variants of unknown functional consequence (26). Positioned at this interface between clinical genetics and electrophysiology, novel computational tools may be helpful to generate supporting evidence. Previously, Heyne et al. demonstrated the use of a machine learning method, gradient boosting, to predict the overall functional effect of missense variants in voltage-gated sodium or calcium channels (27). They used known genotype-phenotype correlations to infer likely functional labels for supervised learning, which may not make full use of available clinical information and risks introducing bias and uncertainty to the training data set. In this study, we extend our own previous work on kernel-based multi-task learning methods applied to potassium channel variants (28) by designing a framework capable of integrating functional, structural and clinical data while leveraging prior knowledge on channel family structure.

## Methods

### Data

Recently, Brunklaus et al. conducted a systematic literature search of variants in voltage-gated sodium channels that were functionally assessed in whole-cell patch-clamp recordings of mammalian cells (29). We refer to their Supplementary Methods for the criteria used to assign functional labels to each variant. Of 437 variants, 62 variants were similar to wildtype function or did not have a clear overall gain- or loss-of-function. We removed these variants from further analysis. Our training data set included 375 unique non-synonymous missense variants across the 9 voltage-gated sodium channels (Na_V_1.1-Na_V_1.9). 164 variants were labelled as GOF, 211 variants were labelled as LOF.

### Features

Each variant was identified by its channel, sequence position, and amino acid substitution. Sequence- and structure-based features were extracted and pre-processed as previously described (28). These 62 features included amino acid physicochemical properties, radicality of substitution, paralog and ortholog conservation, secondary and tertiary structure, residue accessibility, disordered regions, and protein-protein interaction or binding sites. We later attempted to include a further two recently described sequence- and structure-based features: i) 3D residue to pore axis distance, which was shown to be independently associated with several clinical features and overall functional effects (30); ii) parsel, a paralog-specific conservation metric (ancestry conditional site-specific selection) (27). Including these features failed to improve model performance (data not shown), and they were discarded from the final model pipeline.

Variant-level information on primary disease and reference for phenotypes were extracted from the training data set and mapped onto Phenotype MIM numbers of the Online Mendelian Inheritance in Man (OMIM) (31). Each OMIM ID is associated with a set of human phenotype ontology (HPO) terms which represent a standardized vocabulary of phenotypic features in human disease (32). These mappings were requested via the HPO REST API (last accessed 22/June/2022 13:35 pm, release v2022-06-11) and resulted in an average of 9 HPO terms per variant (median 9, range 3 – 42 terms). All parents of the HPO terms in the directed acyclic graph from the parent ← child relationships were added to the respective HPO term sets, which resulted in 446 unique terms across the set of all variants, with each variant enriched to an average of 40 terms (median 38, range 12 – 226). Ontology analysis was performed with the ontologyX suite (33).

We considered three different measures for semantic similarity analysis. For the Jaccard index, we take the ratio of the number of elements in the intersection and the number of elements in the union of the two variant sets. A more rigorous definition and proof of positive definiteness of the function has been provided by Gärtner et al. (34). For the Resnik measure, the similarity between two variants is given as the information content (IC) of their most informative common ancestor (MICA) term (35). The IC is defined as − log_2_(*f*) where *f* is the term frequency over all sets and the MICA is the common ancestor term with the highest IC. Lastly, the Lin measure extends the Resnik measure by incorporating the IC of the individual terms (36). Both Resnik and Lin measures do not lead to positive semi-definite (PSD) similarity matrices in ontologies with multiple ancestors such as the HPO, where terms may have more than one parent term each. Before proceeding with kernel-based learning, we approximated the nearest PSD matrix with the methods suggested by Chen et al. (37).

### Models

When we consider phenotypic semantic similarity analysis as a kernel function and assert its positive semidefinite properties, we satisfy Mercer’s condition and can make implicit use of the reproducing kernel Hilbert space (RKHS) induced by our so-called phenotype kernel. We can avoid explicit representation of each variant in a Euclidean space and instead consider only their inner product, also thought of as pairwise similarity in the high-dimensional phenotypic feature space. This similarity-based learning represents a natural extension of semantic similarity analysis towards kernel-based machine learning methods.

For each variant, the vector of sequence- and structure-based features is transformed into pairwise similarities with a radial basis function (RBF) kernel *K*_*n*_ with 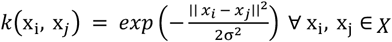 (38). Task similarity, i.e. the hierarchical taxonomy of voltage-gated sodium channels across the channel superfamily, is represented as an alignment-based tree distance measure and enables multi-task (parallel transfer) learning. Both methods have been previously described (28). We can consider each of these three data input types of our variants (channel, sequence/structure, and phenotype) as different measurement modalities with individual kernel matrices for each modality. First, each kernel matrix is normalized in feature space:

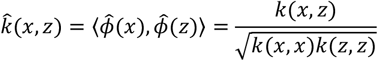

Multiple kernel learning then allows us to represent these modalities in a single kernel matrix *K*, either by assigning uniform weights or by learning a linear combination of matrices *K*_*m*_ with weights *β*_*m*_ such that:

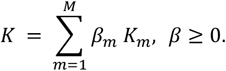

Global weights assign the same weight over the whole input space, which may not be desirable. Several methods of localized multiple kernel learning have instead been proposed (39). We implemented hierarchical decomposition multi-task multi-kernel learning (MTMKL) by Widmer et al. (40). This method makes use of prior knowledge in the form of a tree structure *G*, which in our setting is the taxonomy that relates voltage-gated sodium channels, where each channel is a task *t*_*i*_ corresponding to a leaf in *G*. Given the set of tasks *T* = {*t*_1_, *t*_2_, …, *t*_*T*_}, let the subsets of tasks *S*_*i*_ ⊆ *T* be the leaves of the sub-tree rooted at each node *n* of *G*, that is *leaves*(*n*) = {*l*|*l* is descendant of *n*} (40). The following definition for the kernel function is given:

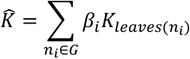

A visualization of these kernel matrices is shown in Supplementary Figure 1. This method also offers additional insight (40). First, the localized kernel weights of each task allow us to see which modality is important for which task. Secondly, we can refine our prior task similarity by learning latent task similarity from the data. If two tasks *t*_*k*_ and *t*_*l*_ are present in subsets *S*_*i*_ that are assigned weights *β*_*i*_, we can define their latent similarity by summing up the weights of their shared task sets where the set of tasks sets containing task *t*_*k*_ is defined as *T*_*k*_ = {*S*|*t*_*k*_ ∈ *S* ∧ *S* ∈ *T*}, where *T* is the set of all tasks, such that the pairwise similarity of tasks *γ*_*k,l*_ is given as:

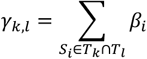

We trained a support-vector machine (C-SVM) on the resulting kernel matrix. Hyperparameter tuning was performed via grid search across cost C = {1e − 4, 1e − 2, …, 1e4}, the RBF kernel hyperparameter σ = {1e − 5, 1e − 4, …, 1e0}, and hierarchical multi-task learning baseline similarity parameter *a* = {1, 3, 5, 10, 100}. The choice of semantic similarity measure and PSD matrix approximation were included in the set of hyperparameters. Hyperparameter tuning and model assessment were done with nested *k*-fold cross validation, *k* = 5, where the model is trained and tested on different random sub-partitions of the data throughout multiple exhaustive iterations to obtain an estimate of the generalization performance. For illustrative purposes, a pseudocode and graphical representation of *k*-fold cross validation is provided in our previous work (28). We report the following metrics: i) Accuracy, the number of correct classifications divided by the number of all classifications; ii) Sensitivity, or true positive rate (TPR); iii) Specificity, or true negative rate (TNR); iv) Matthews correlation coefficient (MCC), a symmetric correlation coefficient between true and predicted classes; iii) Cohen’s Kappa (*κ*), a measure of agreement between true and predicted classes; iv) F_1_ score, the harmonic mean of precision and recall; v) The area under the receiver operator characteristics (AU-ROC) curve; vi) and the area under the precision-recall curve (AU-PRC). Further model information is available in our AIMe Registry report (aime.report/xmSKHj).

### Validation

For baseline comparison, we trained one SVM on variants from each task (channel-specific model, Dirac) and one SVM on variants from all tasks (global model, Union). For internal validation, we implemented gradient boosting as described by Heyne et al. (27), training and testing their method on the same splits of our data set. We also implemented the decision rule model proposed by Brunklaus et al., assigning labels based on missense pairs of identical paralog variants across the sodium channel family (29). For external validation, we conducted a systematic literature search of PubMed with the search string (((SCN1A) OR (SCN2A) OR (SCN3A) OR (SCN4A) OR (SCN5A) OR (SCN8A) OR (SCN9A) OR (SCN10A) OR (SCN11A)) AND ((gain-of-function) OR (loss-of-function)) OR ((GOF) OR (LOF)) AND ((electrophysiology) OR (patch clamp) OR (voltage clamp))) AND ((“2021/04/28”[Date - Publication] : “2022/07/13”[Date - Publication])). This query captures all publications after the last update of the training data set. 55 studies were identified and screened for missense variants in voltage-gated sodium channels with experimental evidence of whole-cell patch-clamp recordings that yielded an overall gain- or loss-of-function effect (n = 91). Variants that were present in the training data set, i.e. those known from previous functional experiments, were removed from further analysis (n = 61). Predictions were obtained from our own model and the web interfaces of Heyne et al. (funNCion, http://funncion.broadinstitute.org/), and Brunklaus et al. (SCN viewer, https://scn-viewer.broadinstitute.org/).

### Software

This study was carried out in the R programming language, version 4.1.0, with RStudio, version 1.4.1106.

## Results

### Learning with phenotypic similarity outperforms both conventional baseline and state-of-the-art models

We first trained naïve baseline models, one channel-specific support-vector machine (SVM) for variants from every single voltage-gated sodium channel (Dirac), and one global SVM for variants from all voltage-gated sodium channels (Union). We then implemented a method from our previous work: Multi-task learning (MTL) SVM embeds explicit prior knowledge on the structural and evolutionary pairwise similarity between ion channels in the feature space induced by our kernel function, which enables the model to assign more weight when learning from variants of similar channels (28). MTL-SVM outperformed the baseline models across all metrics (Table 1a). Finally, our novel multi-task multi-kernel learning model (MT)MKL approach for GOF/LOF prediction was trained with the Jaccard similarity measure and uniform kernel weights. Including phenotypic similarity information in the machine learning model further improves predictive performance (Table 1a, Figure 1). For comparison to current state-of-the-art methods, we trained a gradient boosting machine as used in previous work by Heyne et al. and the decision rule introduced by Brunklaus et al. (27, 29). For internal validation, these methods were trained and tested on the same data splits of our pre-processed training data using the k-fold nested cross-validation procedure described above. For external validation, all three models were used to obtain predictions for a list of novel missense variants in voltage-gated sodium channels (Supplementary Table 1). Results are shown in Table 1b. In both validation procedures, our multi-task multi-kernel learning model performed best.

**Table 1.**
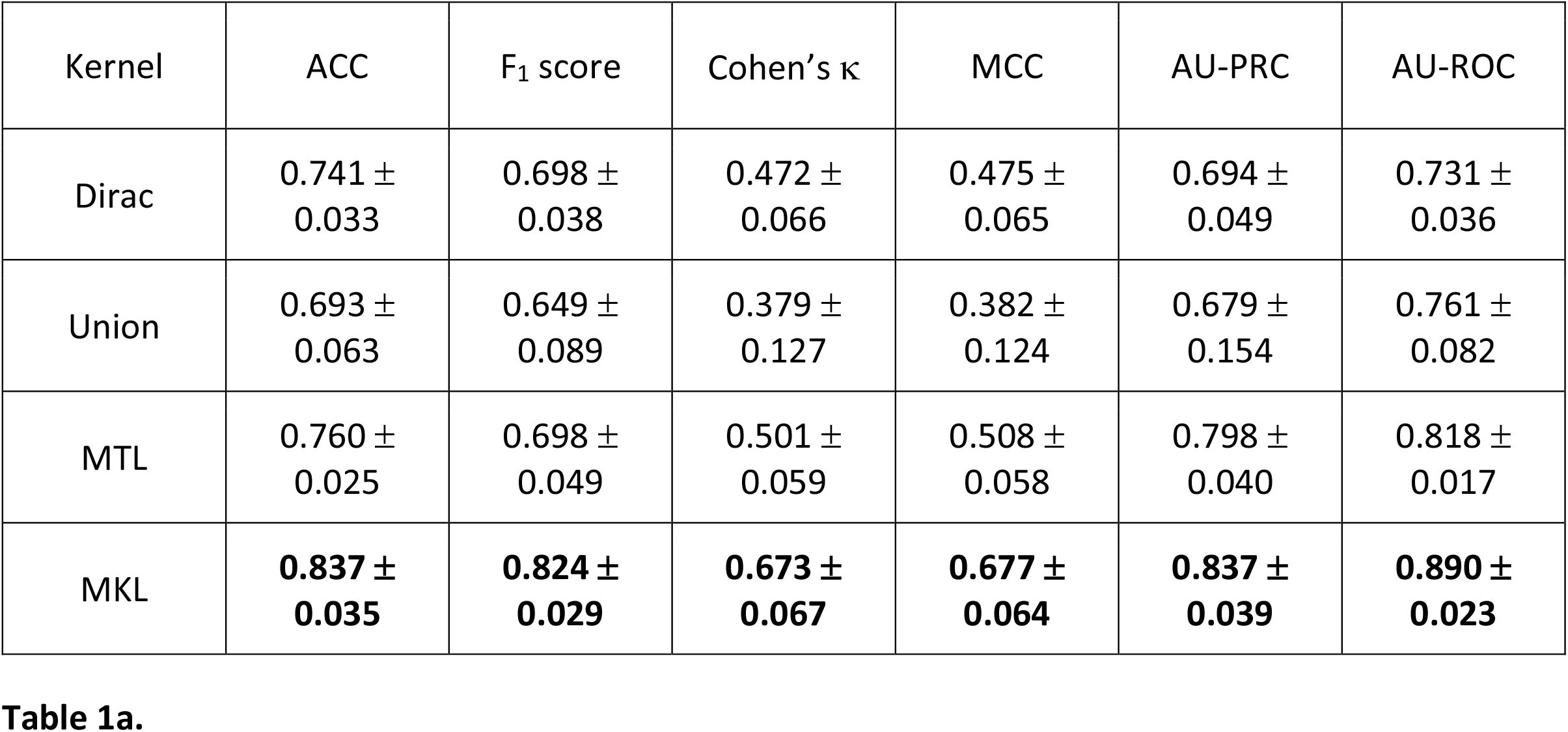
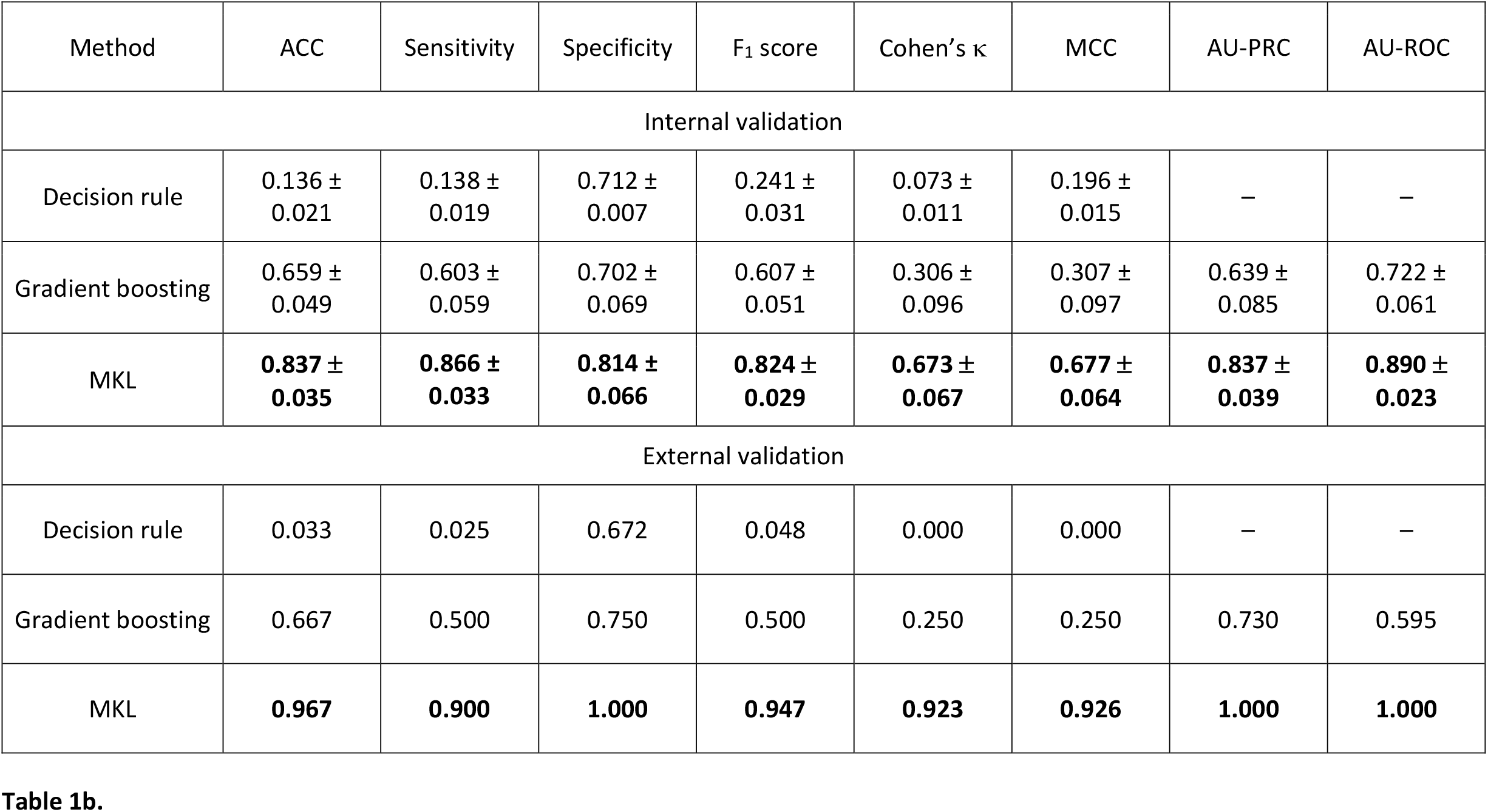
*a*: Nested cross-validation (outer folds) model performance estimates for naïve baseline models (Dirac, Union), our previous state-of-the-art multi-task learning model (MTL), and phenotypic multi-task multi-kernel learning (MKL). Metrics are reported as mean ± standard deviation (SD). *b*: Nested cross-validation performance estimates (internal validation) and external validation on a set of novel missense variants for state-of-the art models and phenotypic multi-task multi-kernel learning (MKL). Note that AU-PRC and AU-ROC are not available for the decision rule as it is not a probabilistic classifier. Phenotypic learning outperforms other models across all metrics and is shown in bold. Abbreviations: ACC – accuracy; AU-PRC – area under the precision recall curve; AU-ROC – area under the receiver operator characteristics curve.

**Figure 1.**
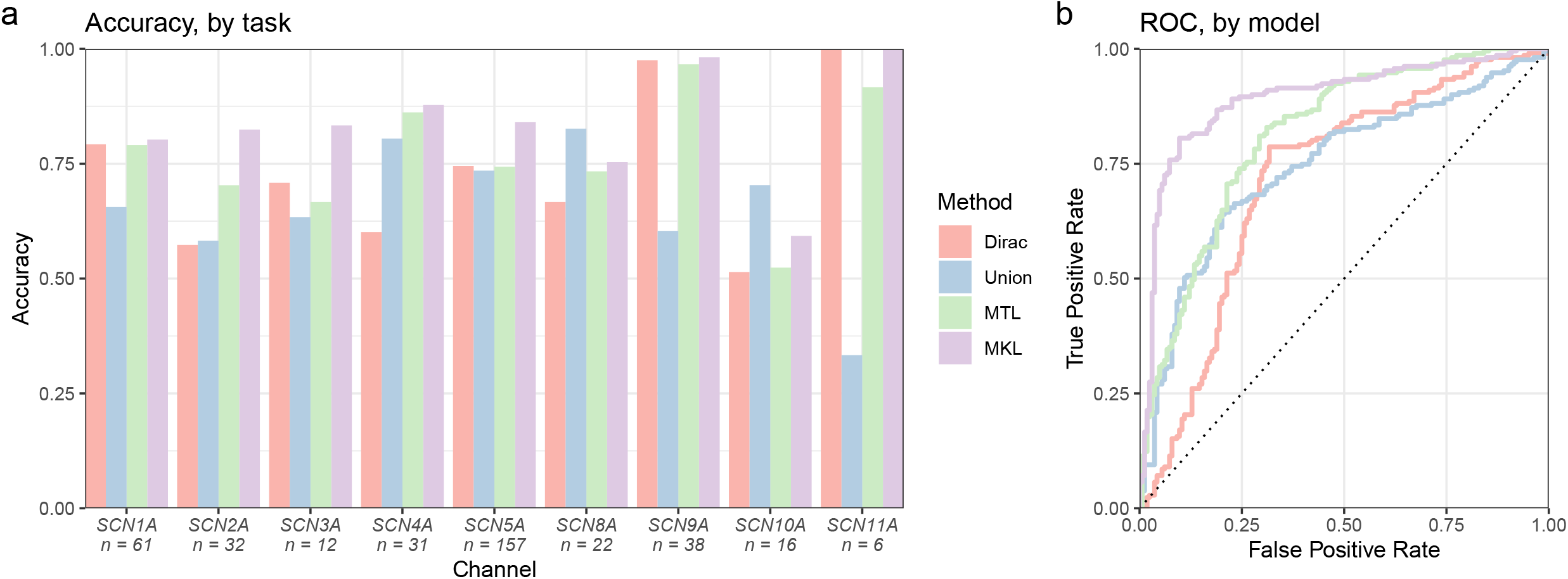
Phenotypic multi-task multi-kernel learning outperforms both conventional and state-of-the-art models. a: Cross-validation model performance estimates of mean accuracy for each task (channel). *b*: ROC curves. The AU-ROC for each model is shown in Table 1. Abbreviations: MTL – multi-task learning; MKL – multi-kernel learning; ROC – receiver operator characteristics.

### Learning with phenotypic similarity is robust across different similarity measures and largely insensitive to phenotypic noise or sparsity

We compared three commonly used similarity measures: Jaccard, Lin, and Resnik. The Jaccard similarity measure always results in a positive semi-definite kernel matrix, which extends naturally to kernel-based learning methods (34). Measures based on information content (IC), such as Lin and Resnik, lead to indefinite kernel matrices for the human phenotype ontology, as it allows for multiple parent terms. This invalidates the underlying theory and would lead to a non-convex SVM optimization problem, which is computationally expensive to solve and is not guaranteed to find the global optimal solution. For similarity-based classification, three methods are available to approximate the nearest positive semi-definite similarity matrix to be used as kernel matrix: spectrum clip, flip and shift (37). The choice of matrix correction method did not have an impact on MKL model performance (Figure 2a, Table 2a).

**Table 2.**
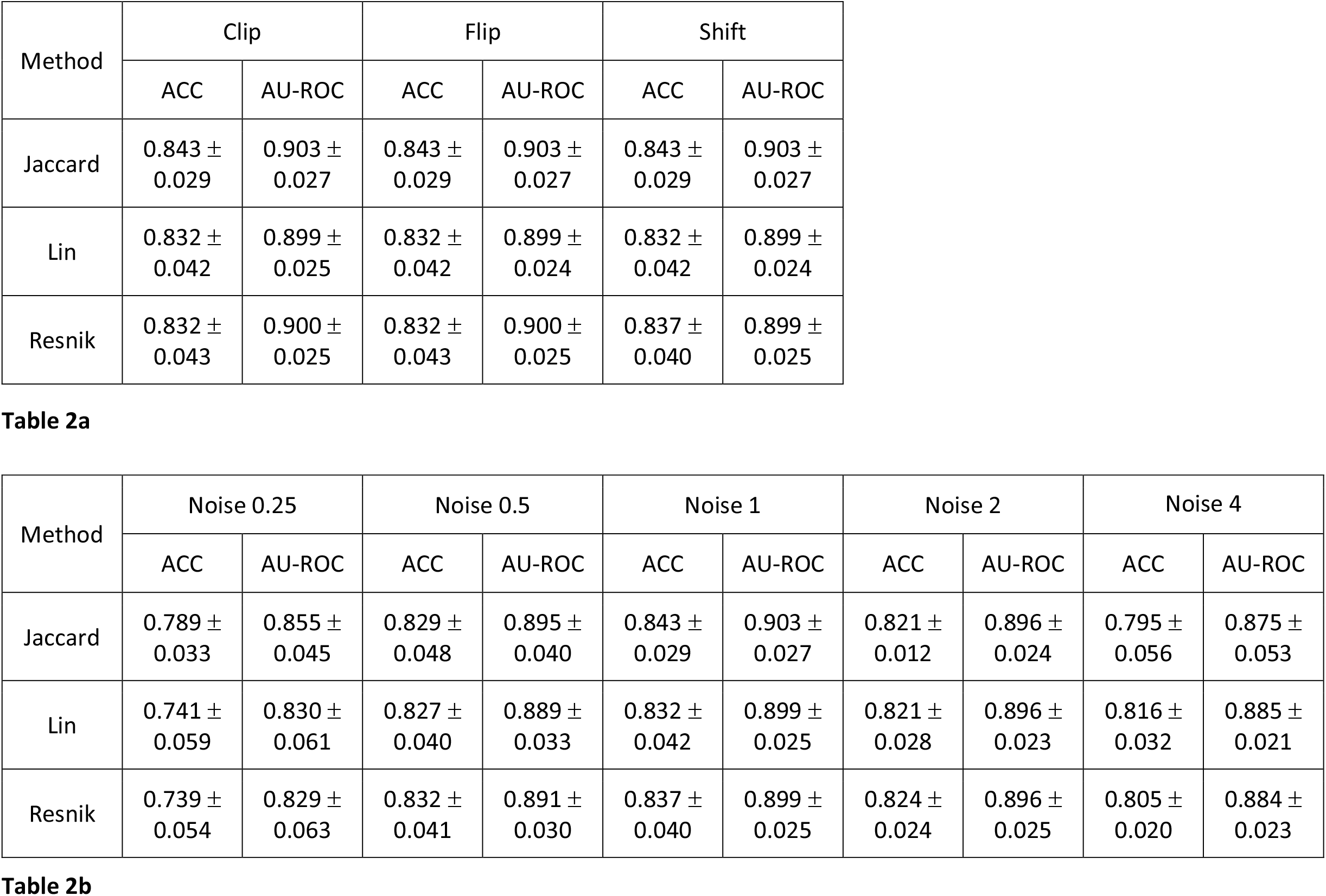
Nested cross-validation (outer folds) model performance estimates. *a*: Model performance of semantic similarity measures over methods of approximating the nearest PSD matrix. The Jaccard index does not require matrix correction but is shown for comparison. *b*: Model performance of semantic similarity measures across levels of simulated phenotypic sparsity and noise (see Figure 2). Abbreviations: ACC – accuracy; AU-ROC – area under the receiver operator characteristics curve.

**Figure 2.**
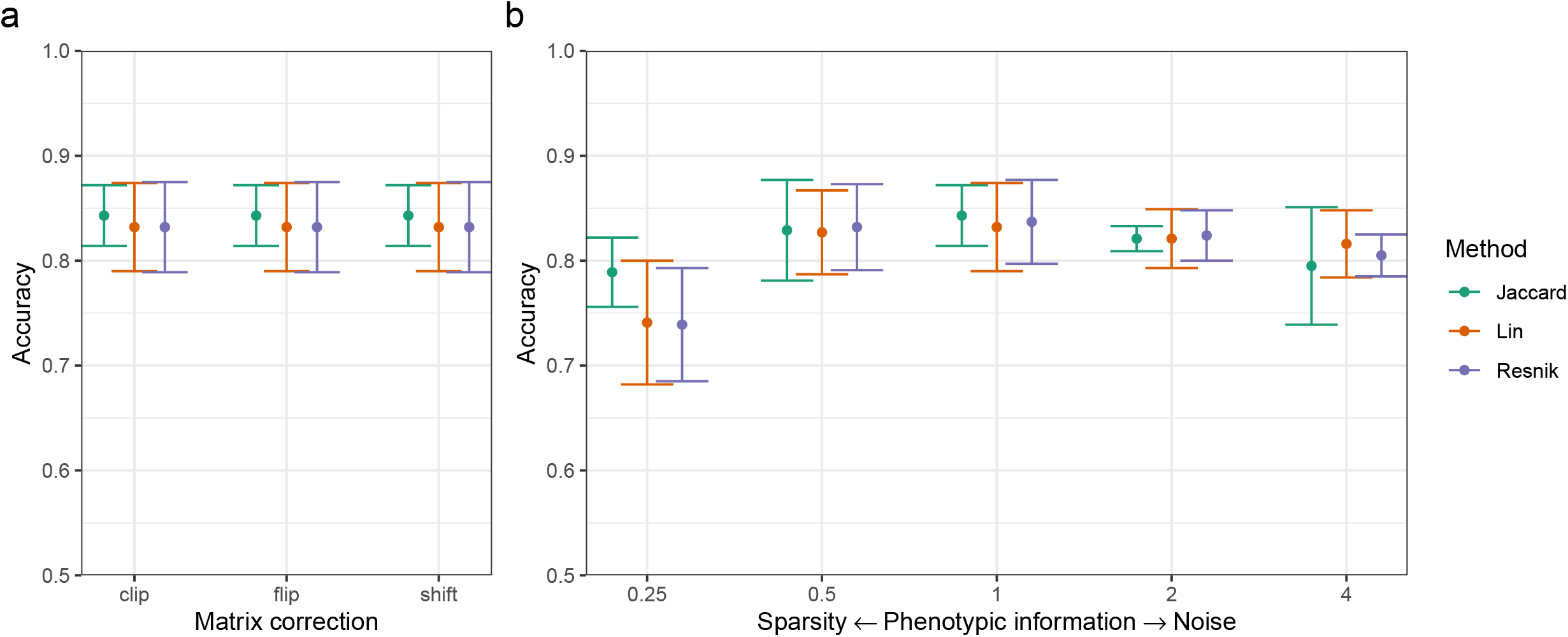
Phenotypic multi-task multi-kernel learning performs well across different similarity measures or matrix correction methods, and is largely insensitive to sparse or noisy phenotypic information. *a*: Dot chart of MKL model mean accuracy and standard deviation for each similarity measure and methods for approximating the nearest PSD matrix. The Jaccard index does not require matrix correction but is shown for comparison. *b*: Dot chart of MKL model mean accuracy and standard deviation for each similarity measure across levels of simulated phenotypic sparsity and noise. Noise is shown as human phenotype ontology term ratio (e.g. noise ratio 0.25 removes 3 out of 4 terms from the set, while noise ratio 4 adds 3 random terms for each original term in the set).

Information content measures are insensitive to varying taxonomic link distances and adapt the static knowledge structure of an ontology to different contexts by including probabilistic estimates (35). We hypothesized that these measures may perform better in imperfectly phenotyped cases. Traditional phenotyping is expensive and prone to misclassification error. Recent studies have introduced ‘silver standard’ phenotyping from data mining of electronic health records and demonstrated advances in automated phenotyping pipelines (41, 42). We simulated sparse phenotypes by removing terms from the set of all terms for each variant at random before computing semantic similarity. As more terms were removed, the information from the phenotype kernel matrix was reduced and the mean accuracy of the MKL model approached that of the MTL model (Figure 2b, Table 2b). Likewise, we simulated noisy phenotypes by sampling (without replacement) additional terms from the set of all possible terms and adding these to the set of terms for each variant before computing semantic similarity. We observed that all semantic similarity measures resulted in models with a stable accuracy even at high levels of simulated noise (Figure 2b, Table 2b).

### Localized multi-task multi-kernel learning with hierarchical decomposition offers biological insight

Previously, we trained the model under the assumption that each kernel matrix (modality) contributes equally by assigning uniform global weights. The global weights for the best convex combination of kernel matrices can also be conveniently learnt from the data with wrapper methods, which have previously been demonstrated as promising tools to work with multi-source genomic data (43). Here, we observed no significant difference in model performance between uniform weights and learnt global weights (Table 3). This is not surprising, as learning weights is most useful when we are working with a large set of kernel matrices and aim to achieve a sparse solution. If we have the prior knowledge to select a small set of adequate kernel matrices, uniform weights are usually sufficient. For theoretical guarantees, see Bach et al. (44).

**Table 3.** Nested cross-validation (outer folds) model performance estimates for each multi-kernel learning method. Uniform weights correspond to the sum of kernel matrices. For global weights, a wrapper method is used to learn the optimal weighted sum of kernel matrices. Hierarchical decomposition learns both localized weights and an optimal weighted sum of subset kernel matrices. The highest mean metric for each method is shown in bold.

| Method | ACC | F<sub>1</sub> score | Cohen's $\kappa$ | MCC | AU-PRC | AU-ROC |
| --- | --- | --- | --- | --- | --- | --- |
| Uniform weights | 0.837 ±<br>0.035 | 0.824 ±<br>0.029 | 0.673 ±<br>0.067 | 0.677 ±<br>0.064 | 0.837 ±<br>0.039 | 0.890 ±<br>0.023 |
| Global weights | 0.843 ±<br>0.038 | 0.832 ±<br>0.029 | 0.684 ±<br>0.071 | 0.691 ±<br>0.069 | **0.880 ±**<br>**0.046** | **0.915 ±**<br>**0.018** |
| Hierarchical<br>decomposition | **0.853 ±**<br>**0.016** | **0.838 ±**<br>**0.028** | **0.705 ±**<br>**0.031** | **0.715 ±**<br>**0.018** | 0.858 ±<br>0.058 | 0.912 ±<br>0.025 |

Global weights assign the same weight to a kernel for all pairs of samples. Therefore, localized weights for each task in the input space may be more appropriate for precision medicine approaches (39). We implemented hierarchical decomposition multi-task multi-kernel learning, which makes efficient use of our prior knowledge on channel taxonomy to learn localized weights for each task and each subset of tasks that are leaves of subtrees rooted at each node (40). Thus, we could identify tasks and subsets of related tasks where the weight of our phenotypic similarity kernel was high. These tasks were likely to contain phenotypic information that contributed towards classification between GOF and LOF, which may be useful for downstream analysis. A dendrogram of voltage-gated sodium channels and their local weights is shown in Supplementary Figure 2. The model correctly identified channels with known or shared genotype-phenotype correlations, e.g. *SCN1A, SCN2A, SCN3A*, and *SCN5A*. A weighted linear combination of these subset kernel matrices was then used as a composite kernel matrix for the model, where the weights were learnt with wrapper methods. Here, the weight vector favoured the kernel matrix of the subset of all tasks (root, wt = 0.93). Thus, the model recognized that voltage-gated sodium channels were similar enough to be learnt together. Overall, hierarchical decomposition offered a modest improvement in model performance (Table 3).

### Finding difficult-to-predict variants in the reproducing kernel Hilbert space

Kernel-SVMs are considered ‘black box’ machine learning models. However, we may still be able to indirectly gather insight on model decisions. Figure 3a shows the histogram of projections for our model, as introduced by Cherkassky et al. (45). Training data were projected onto the normal vector and plotted by their feature space distance to the optimally separating hyperplane. Samples within the range {*x* ∈ ℝ: −1 ≤ *x* ≤ 1} were support vectors. If a sample is close to the hyperplane, model prediction confidence decreases. The set of samples within an arbitrary distance of {*x* ∈ ℝ: −0.5 ≤ *x* ≤ 0.5} (“low confidence”) was compared to the set of samples outside this range (“high confidence”) by feature correlation with an unpaired two-samples Wilcoxon test. We adjusted p-values for multiple testing with the Benjamini-Hochberg method. Features that were significantly (*α =* 0.05) associated with variants that were hard to predict (“low confidence”) are shown in Figure 3b and Figure 3c. This included variants in hotspots along the channel family sequence alignment corresponding to repeat domain IV, as well as variants with a high accessible surface area. For the final model, “low confidence” variants (n = 171) were predicted with a mean accuracy of 0.760, while “high confidence” variants (n = 204) were predicted with a mean accuracy of 0.931.

**Figure 3.**
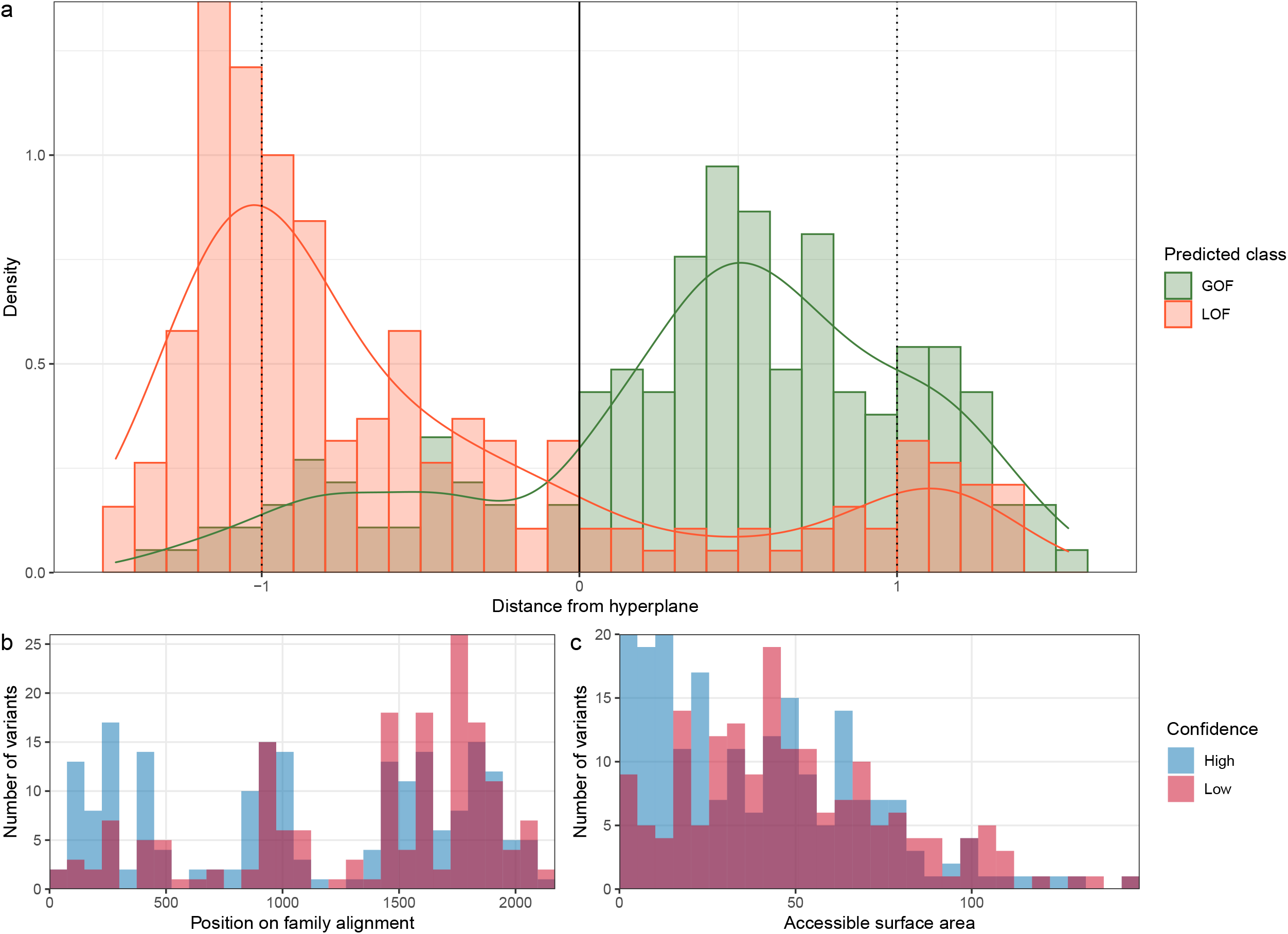
*a*: Cherkassky histogram of projections for the phenotypic multi-task multi-kernel learning model. *b-c*: Features that were significantly different between variants predicted with low confidence (distance to hyperplane ≤ 0.5) and high confidence (distance to hyperplane ≥ 0.5). The accessible surface area feature is based on surrogate predictions by NetSurfP-2.0 and is given in Å (59).

## Discussion

Learning with phenotypic similarity as features represents a natural extension of previous sequence- and structure-based machine learning algorithms (27, 28). Previous studies have demonstrated that clinical features represent an important source of information that may be sufficient to clearly delineate between GOF and LOF in several channelopathies (46, 47). Here, we introduce the concept of extending semantic similarity analysis of human phenotype ontology (HPO) terms towards kernel-based machine learning methods. Beyond the gain in predictive performance, this method has several distinct advantages: i) HPO terms have become the standardized language in clinical genetics. They have been used to structure phenotypic information in large-scale sequencing consortia (42), enable deep phenotyping at the point of care (48), and drive phenotype matching software such as Exomiser (32); ii) Conventionally, all features used for model training must be available at time of prediction. That is, presence or absence of all clinical information would have to be explicitly coded which quickly becomes intractable for complex models. The underlying knowledge graph of the ontology and use of similarity as features solves this issue; iii) Likewise, genotype-phenotype correlations do not have to be explicitly provided to the model but are instead learnt from the data. Identifying channels with high phenotypic similarity matrix weight offers insight for downstream analysis.

Extending our previous multi-task learning framework with multiple kernels allows us to include more information in the feature space. As the kernel matrix represents the information bottleneck for the analysis method (49), including more information has recently contributed towards data integration in multi-omics approaches (50, 51). Accordingly, we observed a large gain in performance for our model when compared to conventional baseline and current state-of-the-art methods. Model performance was robust across different semantic similarity measures. For the final model, we chose the Jaccard index as it represents a computationally efficient method that is guaranteed to result in a similarity matrix that fulfils model assumptions, avoiding theoretical consistency issues that may arise with the Lin and Resnik measures (37). Encouragingly, model performance remained largely stable across simulated sparse and noisy phenotypes which more closely reflects the reality of routinely collected health data such as electronic health records and heterogeneous patient cohorts.

Localized multi-kernel learning by hierarchical decomposition offered insight into implicit phenotype correlations, identifying channels where clinical information is of additional use. The method is also capable of refining prior knowledge on task similarity with latent task similarity learnt from the data. Model performance only increased modestly when compared to the global model as the tasks were overall similar to each other. While this is true for the family of voltage-gated sodium channels (52, 53), this general framework is a promising approach that we expect to scale well to larger task structures (such as unified models of different ion channel families) and to taxonomies with unequal edge lengths (40). Lastly, feature correlation from the histogram of projections of our model identified two features that are associated with hard-to-predict variants: Location in repeat domain IV and a high residue accessible surface area. This information reflects our current mechanistic and structural understanding of sodium channels (52), and helps users identify whether their prediction is in a low-confidence region. That said, localized multi-kernel learning failed to accurately identify some channels with known genotype-phenotype correlations. For *SCN4A*, the clinical spectrum of myopathies is well known (1, 54). For *SCN8A*, recent work by Johannesen et al. has elegantly demonstrated that age at seizure onset and type of epilepsy can delineate between functional effects (24). It is likely that our model currently does not have the resolution necessary to accurately discriminate based on detailed concepts such as myopathy subtypes or age at seizure onset, as the training data set contained only coarse key phenotypes. Additionally, *SCN9A* was learnt with low phenotypic weight likely due to class imbalance as the training data only included a single *SCN9A*-LOF. Phenotypic learning will benefit from large deeply phenotyped cohorts and more granular representation of concepts within the ontology (42).

While our model represents a large step forward in functional variant prediction, several limitations still apply. Our model predicts functional effect at the alpha subunit level, but does not include data from auxiliary beta subunits (*SCN1B-SCN3B*) or other modulator proteins (e.g. *FGF12*) which may themselves affect channel expression levels, gating kinetics, or neuronal network activity (55-57). The model is trained on data from whole-cell patch-clamp recordings of mammalian cells but includes no further information on experimental metadata. Hence, results may differ depending on the choice of expression system (14). Regarding the results of internal and external validation, filtering variants with a clear net gain- or loss-of-function introduces selection bias by neglecting those variants with complex or mixed functional effects. For these variants in particular, electrophysiology remains the gold standard for evaluating functional effects. Most importantly, in silico tools like our model must not be considered strong or causal evidence by the genetic testing standards of the American College of Medical Genetics and Genomics (58), and should not be solely used to establish variant pathogenicity or to directly inform diagnosis and management of affected individuals. These novel tools require further validation and should currently be considered as prediction engines for research use, not as treatment decision support systems.

## Conclusions

Learning with phenotypic similarity represents a novel step from harmonized ontological representations of clinical concepts towards kernel-based machine learning methods. Our framework is robust to phenotypic noise, computationally efficient, scalable to different channelopathies, and outperforms other state-of-the-art methods. The model generates supporting evidence for variant interpretation, and may aid clinicians and biophysical researchers in generating new clinical and experimental hypotheses.

## List of abbreviations

AU-PRC: area under the precision-recall curve
AU-ROC: area under the receiver operator characteristics
BrS: Brugada syndrome
DEE: developmental and epileptic encephalopathy
FHM3: familial hemiplegic migraine type 3
GOF: gain-of-function
HPO: human phenotype ontology
IC: information content
LOF: loss-of-function
LQT3: long QT syndrome type 3
MCC: Matthews correlation coefficient
MICA: most informative common ancestor
MKL: multi-kernel learning
MTL: multi-task learning
MTMKL: multi-task multi-kernel learning
NDEEMA: neonatal DEE with movement disorder and arthrogryposis
OMIM: Online Mendelian Inheritance in Man
PEPD: paroxysmal extreme pain disorder
PSD: positive semi-definite
RBF: radial basis function
RKHS: reproducing kernel Hilbert space
SCB: sodium channel blockers
SCN1A: sodium channel protein type 1 subunit alpha
SeLIE: self-limited (familial) infantile epilepsy
SVM: support-vector machine
TNR: true negative rate
TPR: true positive rate;

## Ethics approval and consent to participate

Not applicable.

## Consent for publication

Not applicable.

## Availability of data and materials

All data and code generated during this study are available on Zenodo (DOI 10.5281/zenodo.7113318). The web-based user interface is available on GitHub (github.com/christianbosselmann/SCION).

## Authors’ contributions

Resources: HL, NP. Supervision: HL, NP. Methodology: CMB. Data Validation: UBSH. Software: CMB. Formal analysis: CMB, NP. Writing – Original Draft: CMB. Visualization: CMB. Writing – Review & Editing: UBSH, HL, NP. All authors have read and approved the final version of the manuscript.

## Acknowledgements

Not applicable.

## Supplementary Material

**Supplementary Figure 1.**
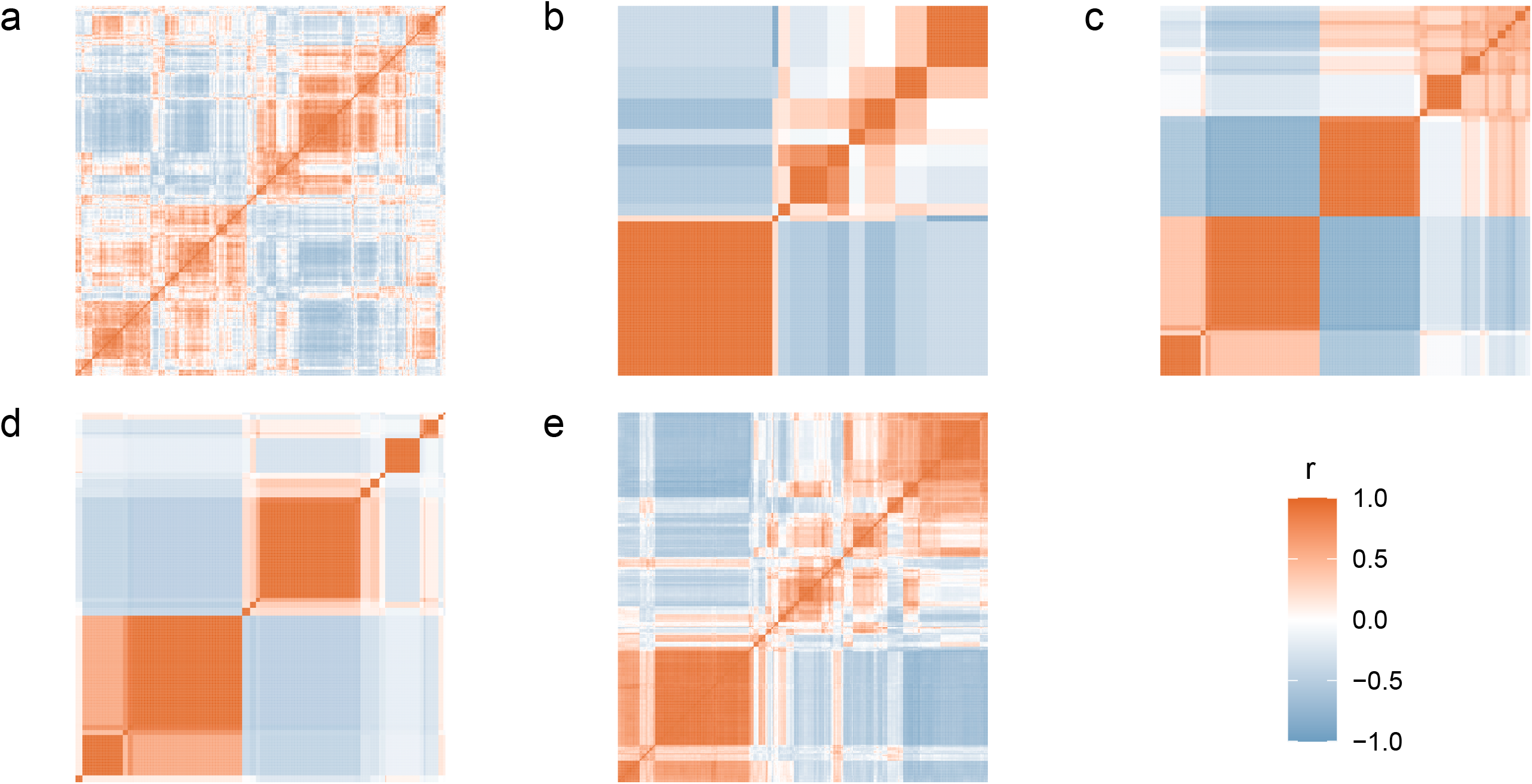
Kernel matrix visualization as correlation plots ordered by hierarchical clustering. *a*: Sequence- and structure-based pairwise similarity matrix. *b*: Task similarity matrix. *c*: Multi-task learning kernel matrix. *d*: Phenotypic similarity matrix. *e*: Multi-task multi-kernel learning matrix.

**Supplementary Figure 2.**
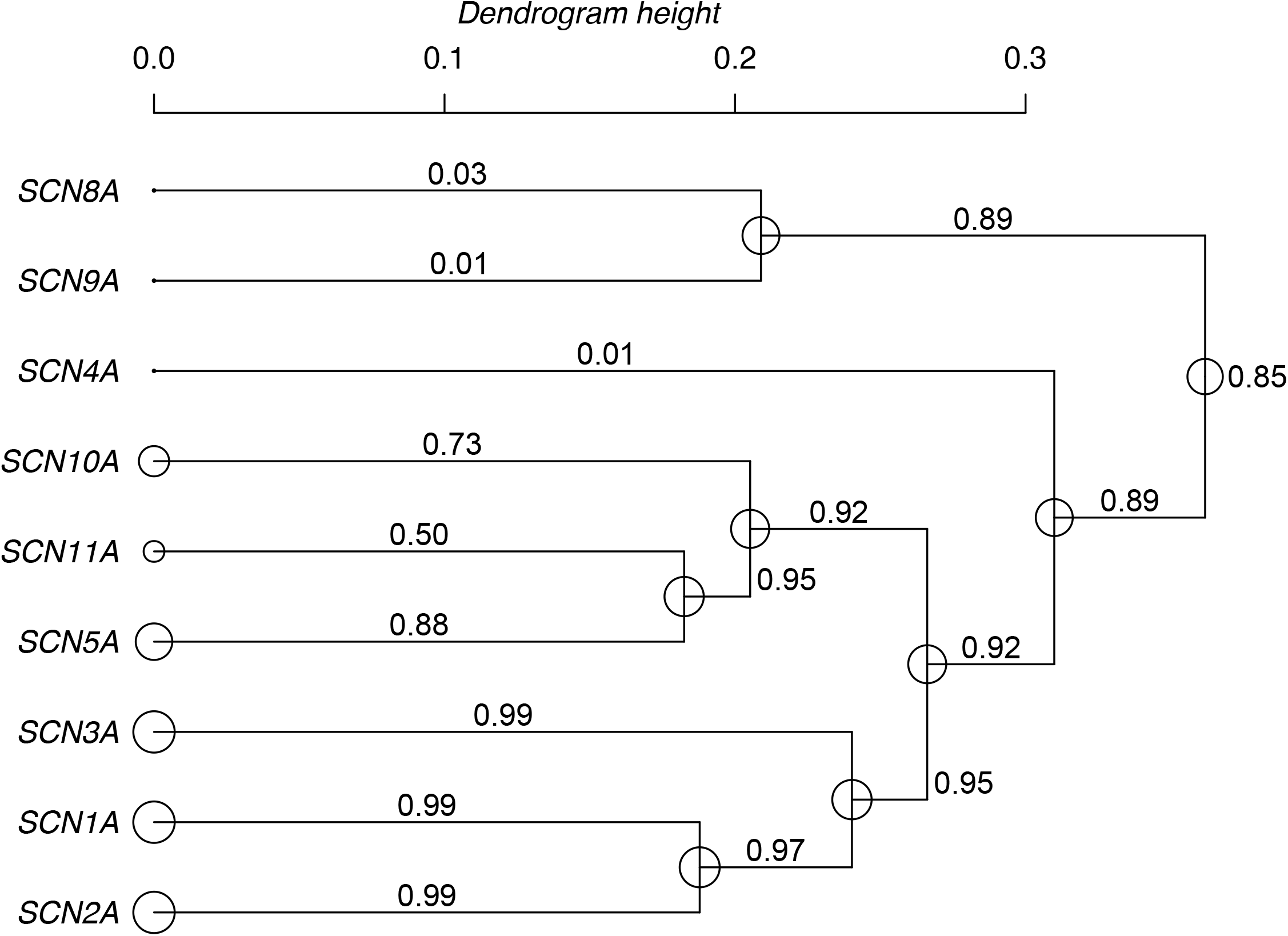
Dendrogram of hierarchical decomposition multi-task multi-kernel learning for voltage-gated sodium channels. Node size and weight labels on the branches correspond to localized phenotypic similarity kernel weights. Uniform weights were assumed for *SCN11A*, as the sample size of the leaf subset was too small. Leaf-to-leaf distance on the dendrogram corresponds to the task similarity used for multi-task learning, with similar tasks clustered together.

**Supplementary Table 1.**
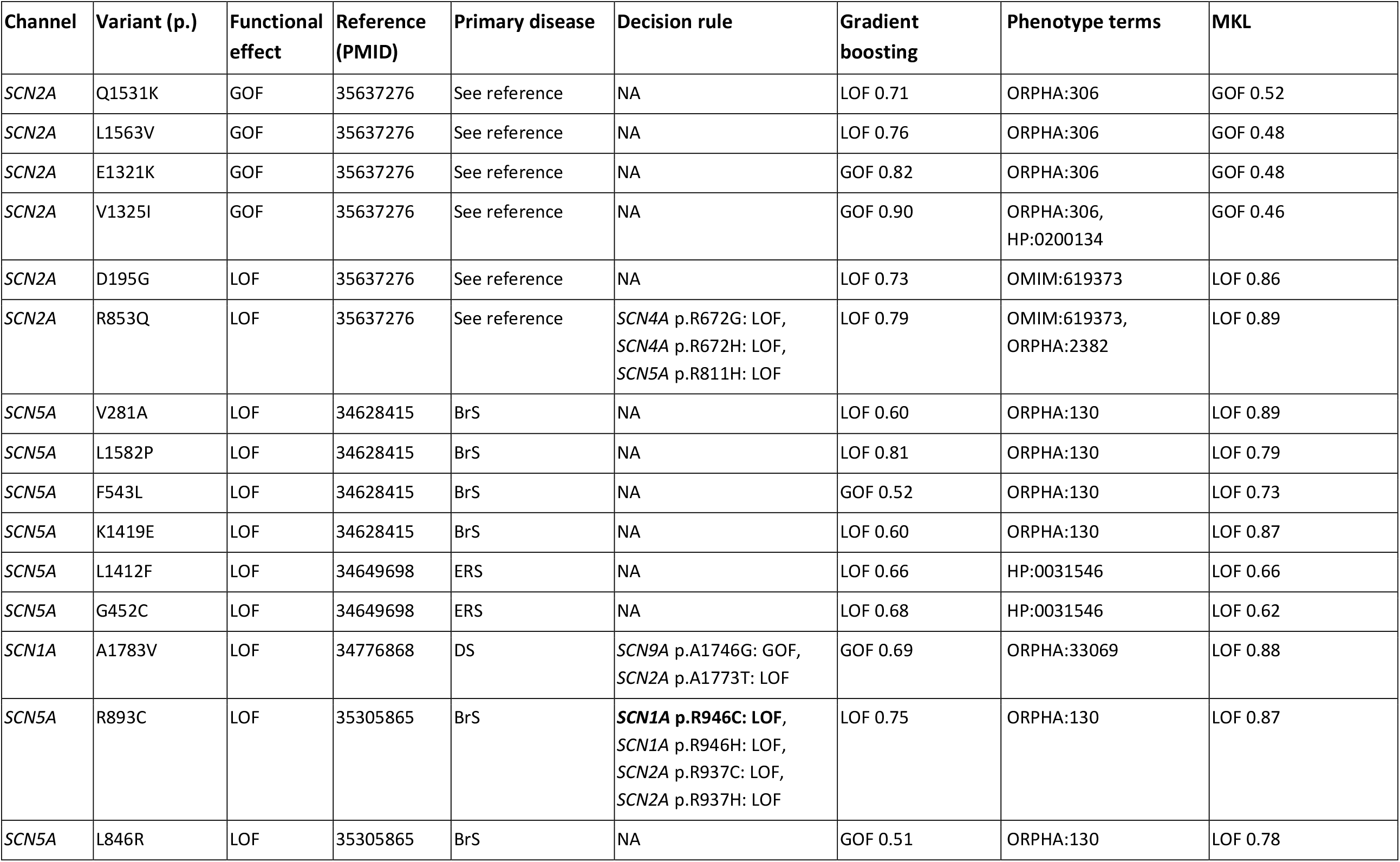
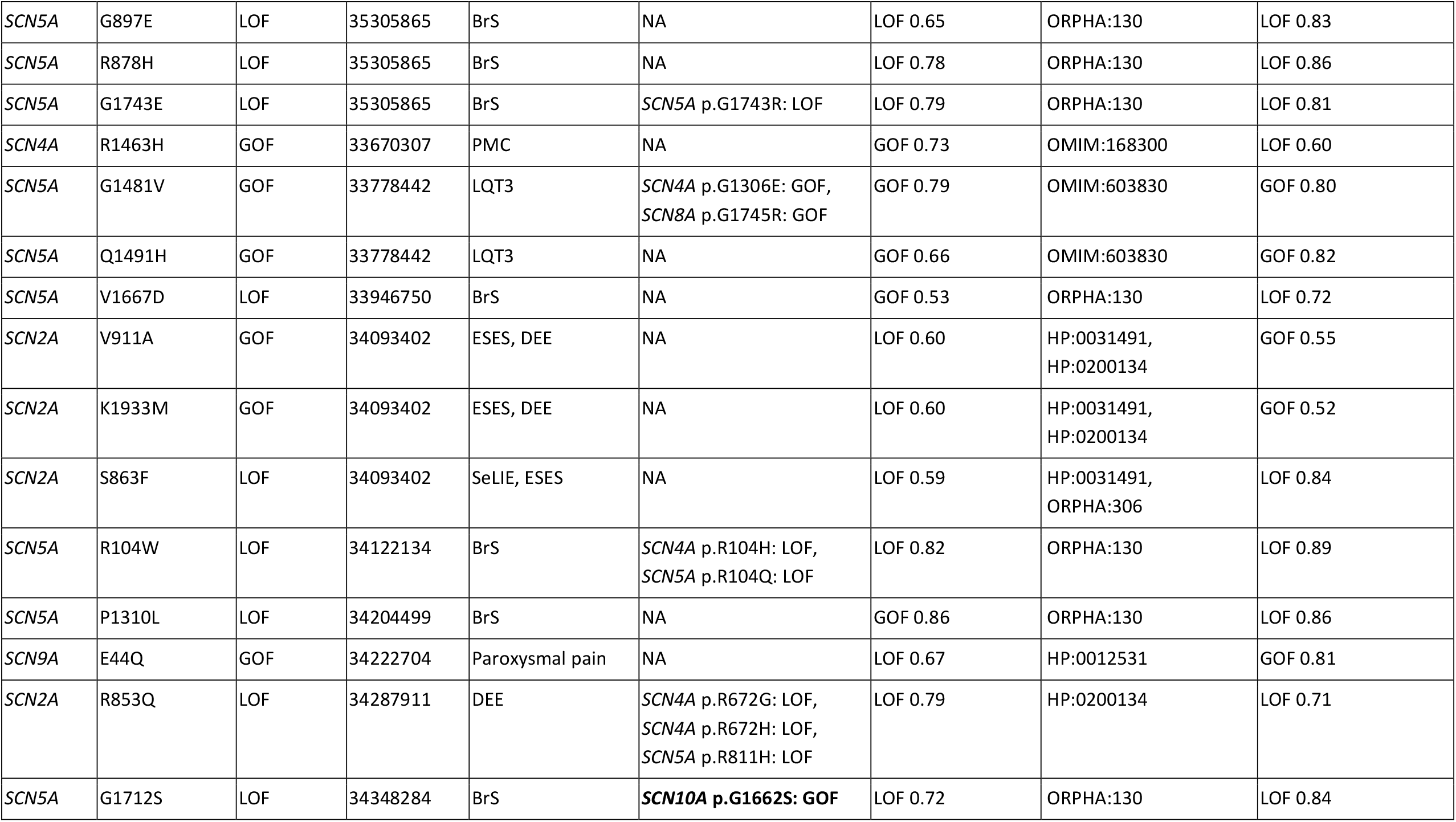
Variants used for external validation. Details on data curation and validation procedure are available in the Methods section. Primary disease corresponds to the core phenotype, while phenotype terms refers to the information provided to the MKL model. For the decision rule, exact matches are shown in bold. Abbreviations: BrS – Brugada syndrome; DEE – developmental and epileptic encephalopathy; DS – Dravet syndrome; ERS – Early repolarization syndrome; ESES - electrical status epilepticus during slow-wave sleep; GOF – gain-of-function; LOF – loss-of-function; LQT3 – long-QT syndrome 3; PMC – paramyotonia congenita; SeLIE – self-limited (familial) infantile epilepsy.

